# Imaging LexA degradation in cells explains regulatory mechanisms and heterogeneity of the SOS response

**DOI:** 10.1101/2020.07.07.191791

**Authors:** Emma C. Jones, Stephan Uphoff

## Abstract

The SOS response functions as the central regulator of DNA repair and mutagenesis in most bacteria and stands as a paradigm of gene networks controlled by a master transcriptional regulator, LexA. We developed a single-molecule imaging approach to directly monitor the LexA repressor inside live *Escherichia coli* cells, demonstrating key mechanisms by which DNA-binding and degradation of LexA regulates the SOS response *in vivo.* Our approach revealed that self-cleavage of LexA occurs frequently during unperturbed growth and causes substantial heterogeneity in LexA abundances across cells. LexA variability underlies SOS gene expression heterogeneity and triggers spontaneous SOS pulses, which enhance bacterial survival in anticipation of stress.

---

Auto-cleavage of the SOS transcriptional repressor LexA induces a wide range of cell functions that are detrimental during optimal growth but are critical for survival and adaptation when bacteria experience stress conditions^1^. Besides activating DNA repair, the SOS response triggers filamentous growth, hypermutation^2^, horizontal gene transfer^3^, antibiotic persistence^4–6^, and the induction of toxins^7^, virulence factors^8^, and prophages^9^. Considering the fitness burden of these functions, it is surprising that the expression of LexA-regulated genes is highly variable across cells^10,7,11^ and that cell subpopulations induce the SOS response spontaneously even in the absence of stress^12–14,6^. Whether this reflects a population survival strategy or a regulatory inaccuracy is unclear, as are the mechanisms underlying SOS heterogeneity. Single-cell measurements of the SOS response have relied entirely on fluorescent gene expression reporters, but their output is itself confounded by noise that is difficult to distinguish from genuine cell-to-cell variation in SOS signalling. Moreover, cellular stress causes global physiological changes that modulate gene expression reporters independently of the SOS response. Although SOS signalling can be measured directly using immunoblots that detect the cleavage of LexA^15^, these assays cannot monitor SOS response dynamics at a single-cell level. As such, it is unknown how heterogeneity in the SOS outputs relates to the underlying input from LexA.

This motivated the development of a microscopy-based approach for measuring DNA-binding and cleavage of LexA directly in individual living cells using single-molecule tracking. We replaced the endogenous *lexA* gene with a HaloTag fusion allowing covalent labelling with the cell-permeable fluorophore Tetramethylrhodamine (TMR) (Fig. 1A). Cells expressing LexA-Halo exhibited normal growth, viability, and SOS gene repression (Fig. S1). SOS induction and survival after ultraviolet light (UV) exposure were partially reduced compared to wild-type cells, but far less so than for SOS-deficient (Ind-) cells expressing a non-cleavable LexA mutant (Fig. S1). These observations, together with the following results, show that the LexA-Halo fusion closely recapitulates native LexA function. High-speed imaging and reversible photoswitching of TMR^16^ allowed tracking of the intracellular movement of hundreds of LexA-Halo molecules per cell (Fig. 1B). During normal growth, cells showed a distinct population of immobile LexA molecules (D ~0.1 μm^2^/s) and a mixture of mobile LexA species with a broad range of diffusion coefficients from D = 0.2 to 12 μm^2^/s (Fig. 1C). The abundance of the immobile population (P_bound_ = 6.3% of tracks) is equivalent to 41 dimers of a pool of 1300 LexA copies per cell^15^, in agreement with the number of LexA dimers required for the repression of ~40 SOS-regulated genes^17^. To test if immobile molecules are indeed DNA-bound, we examined LexAE71K and LexAE45K mutants that have increased DNA-binding affinity *in vitro*^18,19^. Because LexA auto-regulates its own gene, we expressed Halo-tagged LexA mutants from an ectopic pBAD promoter in a Δ*lexA* strain, thus ensuring that expression levels are equal across different variants. As predicted, both LexAE45K and LexAE71K mutants exhibited a larger population of immobile molecules compared to wild-type LexA expressed from the pBAD promoter (Fig. 1D). Both mutants also showed a shift in the diffusion coefficients of the mobile LexA population, likely reflecting a slow-down due to an increase in transient DNA interactions. Next, we added LexA binding sites by transforming cells with pUC19 plasmids carrying promoters that are regulated by LexA, or the constitutive PpolA promoter as a negative control. Indeed, the bound population increased in the presence of additional PsulA sequences that contain one LexA binding site, and further increased for PrecN with three LexA binding sites (Fig. 1E).

**Figure 1.**
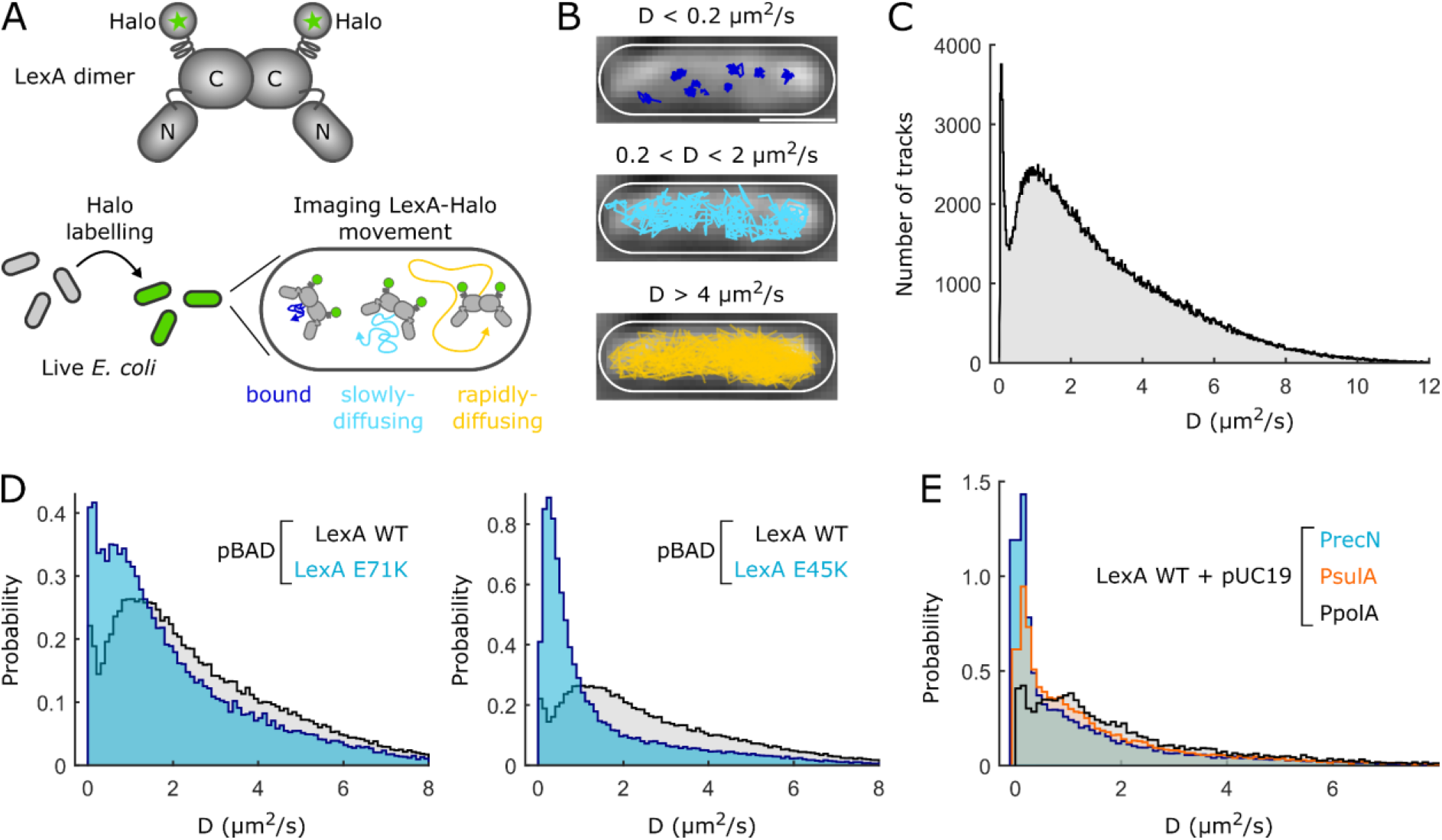
Live-cell single-molecule tracking of LexA repressor. **(A)** Schematic of LexA-Halo fusion protein and imaging procedure. **(B)** Transmitted light image with LexA-Halo tracks coloured according to the diffusion coefficient per molecule in an example cell. **(C)** D distribution of LexA-Halo in untreated wild-type cells (2504 cells, P_bound_ = 6.3%). **(D)** D distributions for pBAD-expressed LexA-Halo mutants with increased DNA-binding affinity, E71K (419 cells, P_bound_ = 10.3%) and E45K (438 cells, P_bound_ = 16.3%), compared to wild-type LexA-Halo (838 cells, P_bound_ = 4.9%). **(E)** D distributions for wild-type LexA-Halo in cells carrying pUC19 plasmids with LexA-regulated promoters PsulA (304 cells, P_bound_ = 18.3%), PrecN (281 cells, P_bound_ = 30.0%), or PpolA control (178 cells, P_bound_ = 10.0%).

The SOS response is triggered when RecA proteins accumulate on single-stranded DNA at damage sites or stalled replication forks. Interaction of LexA with RecA filaments stabilises a conformation that causes LexA to cleave itself^20^, which should separate the Halo-tagged C-terminal dimerization domain (LexA^85-202^-Halo) from the N-terminal DNA-binding domain (LexA^1-84^) in our construct (Fig. 1A). Indeed, LexA-Halo mobility increased following a UV pulse due to a loss of DNA-bound and slowly diffusing molecules accompanied by the gradual appearance of a rapidly-diffusing population (Fig. 2A). LexA self-cleavage exposes recognition motifs for degradation of the protein fragments by Lon and ClpXP proteases^21,22^. The distribution of diffusion coefficients of LexA-Halo 180min post UV exposure was identical to that obtained from untreated cells expressing unconjugated HaloTag (Fig. 2B). Hence, the free HaloTag appears to remain in cells as a rapidly-diffusing species after degradation of the LexA^85-202^ cleavage fragment, consistent with the inability of ClpXP to degrade the HaloTag from its N-terminus^23^. SDS-PAGE and in-gel TMR fluorescence detection confirmed the UV-induced cleavage of the LexA-Halo fusion and its conversion into the free HaloTag (Fig. 2C). Although LexA-Halo diffusion increased after UV exposure in a Δ*clpX*Δ*lon* strain with similar kinetics as in the wild-type, the average diffusion coefficients were lower throughout the response (Fig. 2D). This is consistent with the protease-deficient cells being unable to degrade the cleaved LexA^85-202^-Halo fragment, which has a larger size and thus a lower mobility than the HaloTag alone (Fig. 2D). The relative abundances of the DNA-bound population *(P_bound_*), free LexA pool (*P_free_*), and degraded LexA species (*P_degraded_*) provide a quantitative readout for the progression of the SOS response in live cells (Fig. S2, S3). *P_free_* decayed exponentially with a half-life of 19min after a UV dose of 50J/m^2^, while UV doses of 20 and 5J/m^2^ caused slower and incomplete LexA degradation, as expected^24^ (Fig. 2E, S4).

**Figure 2.**
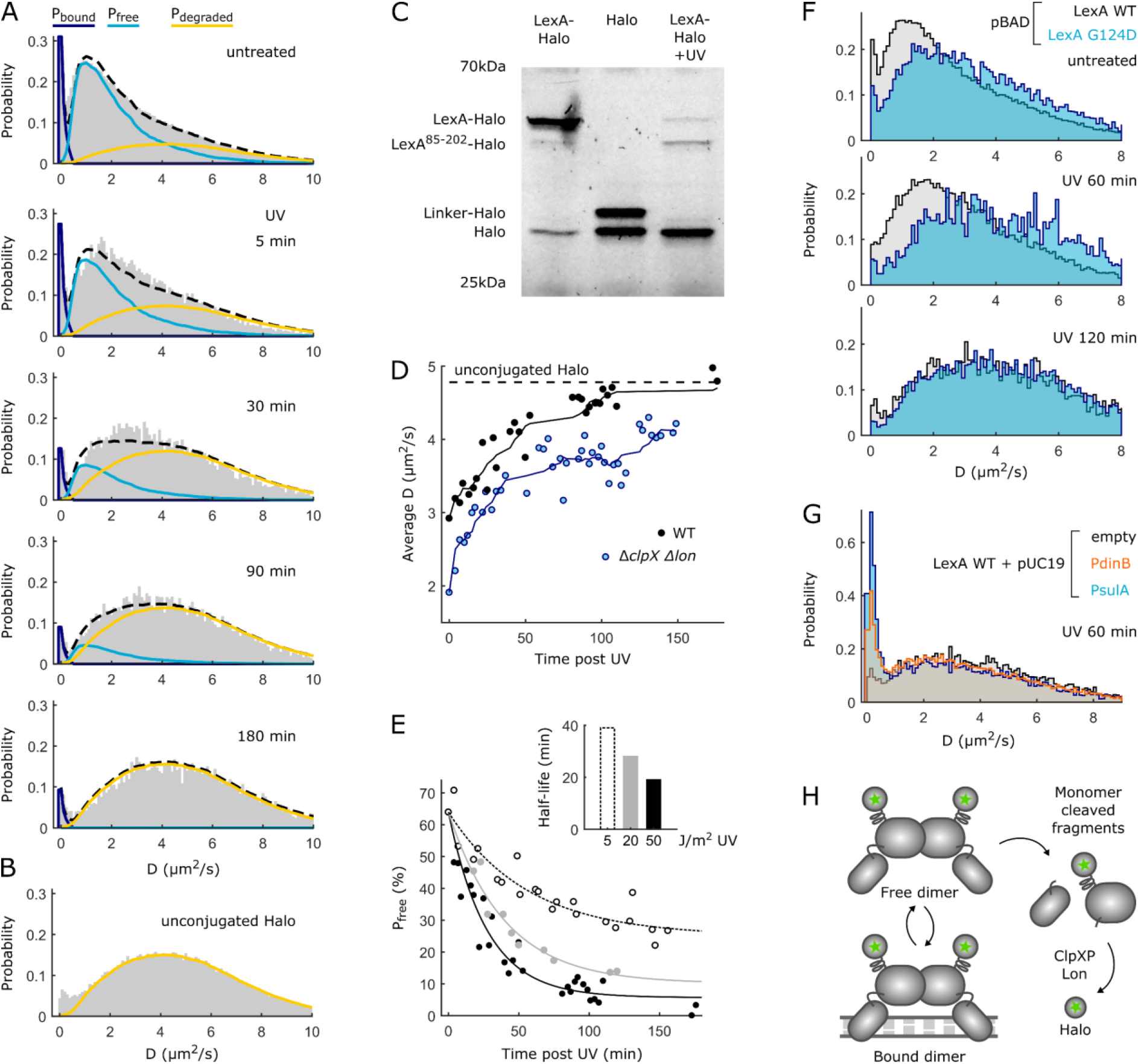
Dynamics of LexA cleavage and degradation in response to DNA damage. **(A)** D distributions for LexA-Halo in untreated cells and at indicated times post 50J/m^2^ UV exposure fitted with mixture model (black dashed lines) of LexA populations P_bound_ (dark blue), P_free_ (light blue), and P_degraded_ (yellow). **(B)** D distribution for unconjugated HaloTag in untreated cells. **(C)** SDS-PAGE in-gel fluorescence from cells expressing LexA-Halo, unconjugated HaloTag, LexA-Halo at 120min post 50J/m^2^ UV exposure. The additional band for the unconjugated HaloTag corresponds to the tag plus the linker. **(D)** ClpXP and Lon degrade cleaved LexA. Average D of LexA-Halo over time after 50J/m^2^ UV in wild-type (black) and Δ*clpX*Δ*lon* strain (blue) with moving average curves. Average D of unconjugated HaloTag in untreated cells shown for reference (dashed line). **(E)** Decay of free LexA pool for different UV doses. Inset: decay half-life from exponential fits. **(F)** LexA is dimeric in cells and monomers are cleaved faster than dimers. D distributions for LexAG124D dimerization mutant compared to wild-type LexA-Halo expressed from pBAD plasmid without treatment (WT: 838 cells, G124D: 173 cells) and post 50J/m^2^ UV at 60min (WT: 257 cells, G124D: 118 cells) and 120min (WT: 197 cells, G124D: 165 cells). **(G)** DNA-binding protects LexA from cleavage. D distributions 60min post 50J/m^2^ UV exposure of cells carrying empty pUC19 plasmid (117 cells, P_bound_ = 2.4%), or pUC19 with PsulA (117 cells, P_bound_ = 12.7%) or PdinB (146 cells, P_bound_ = 7.8%) promoters. (H) Schematic for SOS induction via LexA cleavage and degradation.

To study the role of LexA dimerization, we examined the diffusion profile of the LexAG124D mutant with a 50-fold reduced dimerization constant^25^. It showed a mobile population with increased diffusion coefficient compared to the wild-type (Fig. 2F), matching the expectation that the monomer should be more mobile than the dimer and confirming that LexA is predominately dimeric in cells^20,25–27^. As LexA dimerization is important for promoter recognition^28^, the G124D mutation also diminished the DNA-bound population (Fig. 2F). Self-cleavage of LexA dimers is thought to occur separately for each monomer^25,27,28^. In fact, LexA monomers were not only capable of self-cleavage but the degradation proceeded faster than the wild-type after UV exposure (Fig. 2F).

Although LexA degradation induces an entire gene network, the activation times of different genes follow a specific chronology in response to DNA damage^17,29^. DNA repair genes are activated rapidly, whereas mutagenic DNA polymerases and toxins become induced later in the response as a strategy of last resort. The differential gene induction is attributed to promoter-bound LexA dimers being protected from cleavage^27^, so that the activation time of each gene is governed by the dissociation rate constant of LexA from its promoter. Several lines of evidence from our data confirm this model. First, the decay of the DNA-bound population was slower than the decay of the free LexA pool (Fig. S4). Second, increasing the number of DNA-binding sites in cells delayed LexA degradation (Fig. 2G). Third, degradation was inhibited more strongly in the presence of additional PsulA promoter sequences as compared to PdinB sequences, consistent with the delayed induction of *sulA* compared to *dinB* during the SOS response^17,29^ (Fig. 2G). Together, single-molecule tracking of LexA confirms the key regulatory mechanisms of the SOS response *in vivo* (Fig. 3H).

**Figure 3.**
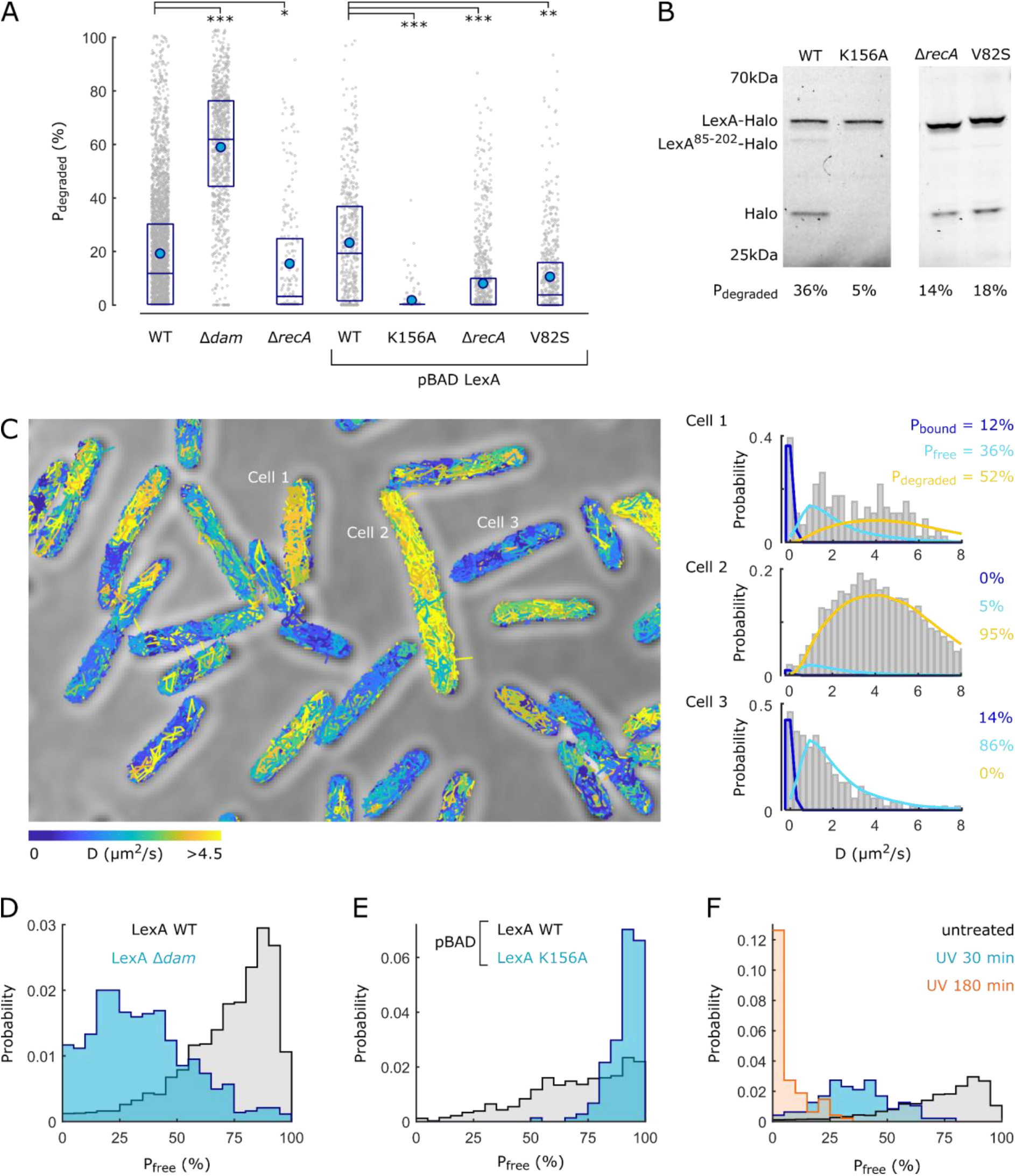
Spontaneous LexA degradation in untreated cells is highly heterogeneous. **(A)** Abundance of degraded LexA-Halo population per cell in wild-type strain (3146 cells), Δ*dam* strain (837 cells), Δ*recA* strain (229 cells). Strains with pBAD expression of wild-type LexA (838 cells), non-cleavable K156 mutant (148 cells), wild-type LexA-Halo in Δ*recA* strain (934 cells), auto-cleavage deficient V82S mutant (582 cells). Boxes: 25-75% percentile, blue dots: averages, lines: medians, grey dots: individual cells, *p<10^-4^, **p<10^-20^, ***p<10^-40^ from two-sided Wilcoxon rank sum test. **(B)** In-gel fluorescence from cells expressing LexA-Halo from pBAD plasmid: wild-type, K156A mutant, wild-type in Δ*recA* strain, V82S mutant. **(C)** Transmitted light image with LexA-Halo tracks coloured according to the diffusion coefficient per molecule. D distributions of three example cells with mixture model for bound, free, and degraded populations. **(D)** Abundance of the free LexA population per cell for wild-type and Δ*dam* strains. **(E)** Abundance of the free LexA population per cell for for wild-type and K156A mutant expressed from pBAD plasmid. **(F)** Abundance of the free LexA population per cell without treatment, and at indicated times post 50J/m^2^ UV.

Quantifying LexA populations in untreated cells via single-molecule tracking or in-gel fluorescence showed that a surprisingly large proportion of LexA molecules become degraded during normal growth (*P_degraded_* = 19.0% or 23.2% for endogenous or pBAD-expressed LexA-Halo) (Fig. 3A-B). Bona-fide LexA cleavage is responsible for this, as spontaneous degradation was diminished in the non-cleavable LexAK156A mutant^25^ (Fig. 3A-B, S5). Perturbed DNA replication and frequent DNA breakage in a Δ*dam* strain^30^ strongly increased LexA degradation (Fig. 3A, S5). However, deletion of RecA did not abolish LexA degradation completely (Fig. 3A, 3B, S5), demonstrating that endogenous DNA damage is not the sole trigger for cleavage during normal growth. In fact, LexA is capable of adopting its auto-cleavable conformation without RecA acting as the co-protease, but the relevance of this pathway inside cells remained uncertain^22,31–34^. We examined the LexA V82S mutation, which blocks the auto-cleavable conformation but allows normal RecA-dependent cleavage^34^. Indeed, LexAV82S still showed basal degradation but at a reduced rate compared to the wild-type (Fig. 3A-B, S5). This, together with our observations for the Δ*recA* mutant, shows that RecA-independent autocleavage contributes to a basal rate of LexA degradation.

Degradation of LexA during normal growth could be the cause of spontaneous inductions of the SOS response associated with the anticipation and survival of sudden stress^6,10–13^. Quantification of LexA populations in single cells revealed an astonishing level of cellular heterogeneity (Fig. 3A & C, Fig. S6), both for endogenous and pBAD-expressed LexA. The broad distributions of *P_degraded_* and *P_free_* show that LexA degradation is triggered very frequently (Fig. 3D), and not solely in a small subpopulation of cells as had been inferred from SOS gene expression reporters^7,11,12^. Evidently, spontaneous SOS induction reflects a continuous scale of LexA degradation levels rather than a distinct regulatory state. Heterogeneity in the free LexA pool was further increased in the Δ*dam* strain, while it was far reduced for cells expressing non-cleavable LexAK156A mutant (Fig. 3E). Despite the variability in the basal state of LexA, UV treatment caused complete and uniform LexA degradation in all cells (Fig. 3F), showing high fidelity in DNA damage signalling.

Our observations raise the question of how LexA’s variability affects SOS gene repression. Single cells growing unperturbed and continuously inside microfluidic channels showed frequent expression pulses of a transcriptional reporter for the SOS response PrecA-GFP (Fig. 4A). The pulse amplitudes had a skewed distribution with many small expression pulses and a long tail of infrequent large pulses (Fig. 4B). These fluctuations were completely absent in cells expressing the non-cleavable LexAG85D mutant (Fig. 4A,C). Furthermore, cells with non-cleavable LexA had an overall lower basal expression level (Fig. 4C), indicating that the frequent spontaneous cleavage in wild-type cells causes partial de-repression of the SOS response. To understand how natural fluctuations in regulatory input levels modulate the transcriptional output of the SOS response, we measured the gene regulation function (GRF)^35^ of LexA at a single-cell level by quantifying LexA-Halo populations and PrecA-GFP expression simultaneously (Fig. 4D). PrecA expression was induced in cells with increased LexA degradation (Fig. 4D-E). Intermediate LexA abundances translated to a continuum of PrecA expression levels (Fig. 4E and inset), explaining how gradual variation in the free LexA pool due to spontaneous degradation creates a continuous scale of SOS expression pulses. This relation was well described by a GRF with an affinity of LexA for PrecA between 2-10 nM^36^ (Fig. 4E). The same GRF also matched PrecA-GFP regulation in the Δ*dam* strain (Fig. 4E), but the distribution of cells along the curve was altered, as expected. Expression of the constitutive PpolA promoter was independent of the state of LexA, confirming that the observed regulation is specific to SOS-controlled genes (Fig. S7).

**Figure 4.**
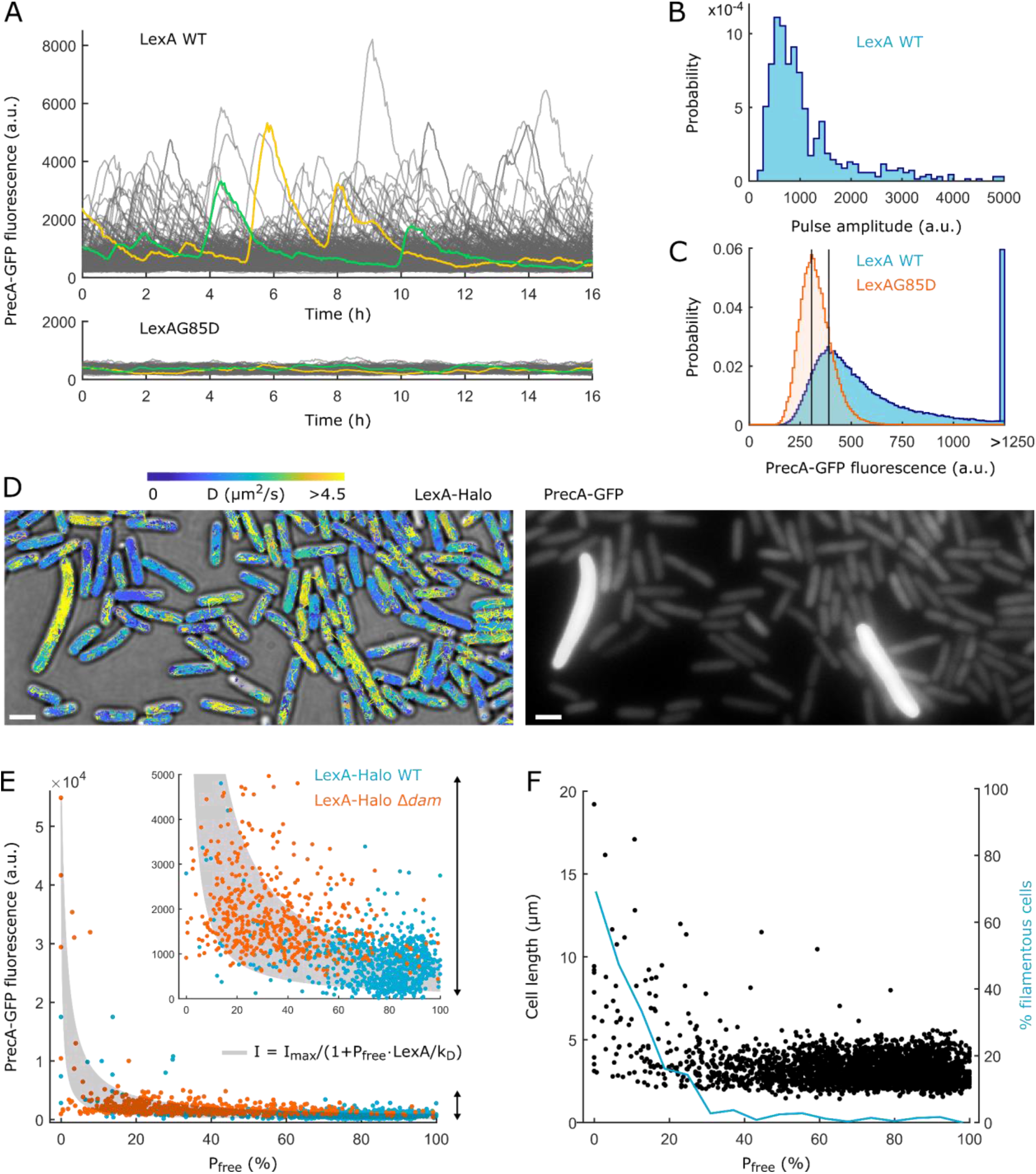
Variability in LexA degradation underlies spontaneous SOS induction and expression heterogeneity. **(A)** Single-cell fluorescence dynamics of SOS expression reporter PrecA-GFP for cells expressing wild-type LexA (474 cells) and non-cleavable LexAG85D (446 cells). Two example cell traces highlighted. **(B)** Distribution of PrecA-GFP pulse amplitudes in wild-type cells. **(C)** Distributions of PrecA-GFP fluorescence for wild-type and LexAG85D cells. Vertical lines show the shift in the mode of the distributions. **(D)** Dual-colour imaging of LexA-Halo tracks (on transmitted light image) and PrecA-GFP fluorescence in the same cells. **(E)** PrecA-GFP fluorescence versus free LexA population per cell for wild-type (blue dots, 853 cells) and Δ*dam* strain (orange dots, 521 cells). Grey area: Gene regulatory function for PrecA-GFP fluorescence (I) as a function of P_free_ (expressed as fraction of molecules [0 1]) with total LexA dimers = 650nM and k_D_ between 2-10nM. Inset: expanded section indicated by arrow. **(F)** Switch-like induction of cell filamentation. Left axis: Cell length versus free LexA abundance per cell (black dots, 2504 cells). Right axis: Percentage of filamentous cells (length > 5μm) across a moving average of free LexA abundance per cell (blue line).

Many of the survival mechanisms induced by the SOS response impose a substantial fitness cost, and therefore must be repressed reliably in the absence of stress, despite the basal LexA fluctuations. SOS induction of the cell division inhibitor SulA is one such mechanism. We found that cells switched to filamentous growth when the free LexA pool declined below a threshold of P_free_<30% (Fig. 4F). Only 5% of cells had such low LexA abundance, and any variation in the free LexA pool above the threshold did not influence cell length or the probability of filamentation (Fig. 4F). The switch-like control of filamentation ensures that cell size homeostasis is robust to LexA variability.

These findings demonstrate that LexA achieves a remarkable balance as a master regulator, ensuring reliable repression and induction of the SOS response, while also generating gene expression variability that can act as a source of cellular differentiation to enhance stress survival of isogenic bacterial populations. Single-molecule tracking of LexA-Halo provides a direct and quantitative readout of the SOS response. The method bypasses key limitations of existing SOS reporters, opening new avenues to understand how the SOS response enables bacterial survival in diverse environments.

## Materials and methods

### Bacterial strains

All strains used in this study are shown in the table below and were derived from *Escherichia coli* AB1157. The LexA-Halo fusion was generated by Lambda Red recombination^37^. We used plasmid pSU007 as a template to insert the HaloTag sequence with a 27 amino acids linker at the C-terminus of the endogenous *lexA* gene followed by a kanamycin resistance cassette. The linker sequence SAEAAAKEAAAKEAAAKEAAAKAAAEF was designed to form an alpha-helical structure. The gene fusion was confirmed by colony PCR and in-gel fluorescence and the allele was moved into AB1157 wild-type strain by P1 phage transduction. The kanamycin resistance gene flanked by frt sites was removed by expressing Flp recombinase from plasmid pCP20. The temperature-sensitive pCP20 was cured by growing cells at 37°C, generating strain SU225.

Strains with chromosomal gene deletions were obtained from the Coli Genetic Stock Center. The alleles were moved into background strain SU225 using P1 phage transduction and selecting for kanamycin resistance. Gene deletions were confirmed by colony PCR. To combine multiple deletions, antibiotic resistance genes flanked by frt sites were removed using pCP20 as above. The Δ*lexA* allele was kindly provided by the Kohli lab (strain SAMP04) and moved into AB1157 Δ*sulA* strain via P1 phage transduction and selection for chloramphenicol resistance. The *sulA* deletion is necessary for viability of Δ*lexA* strains.

For plasmid expression of the LexA-Halo fusion, the *lexA-halo* allele was amplified from SU225 and inserted into plasmid pBAD24 using Gibson assembly kit (NEB) and confirmed by sequencing. LexA point mutants were made using Q5 site-directed mutagenesis kit (NEB) and confirmed by sequencing. Plasmids were transformed into SU315 Δ*lexA ΔsulA* strain and selection of ampicillin resistance, ensuring that the plasmid-expressed LexA-Halo fusion is the only LexA species present.

pUC19 plasmids carrying LexA-regulated promoters were generated by amplifying the promoter sequences from an *E. coli* promoter library ^38^. Insertions were made into the multiple cloning site on pUC19 by restriction digestion using EcoRI and XmaI cut sites. Plasmids were transformed into SU225 expressing chromosomal LexA-Halo.

The PrecA-GFPmut2 and PpolA-GFPmut2 transcriptional reporters were copied from the promoter library plasmids ^38^ and inserted on the chromosome between *tam* and *yneE* genes in the chromosome terminus region ^39^ via Lambda Red recombination and selection for kanamycin resistance. Placement of the transcriptional reporter in the chromosome terminus region ensures it is present at a single copy during most of the replication cycle, thus minimising expression fluctuations due to gene copy number variations.

### List of strains

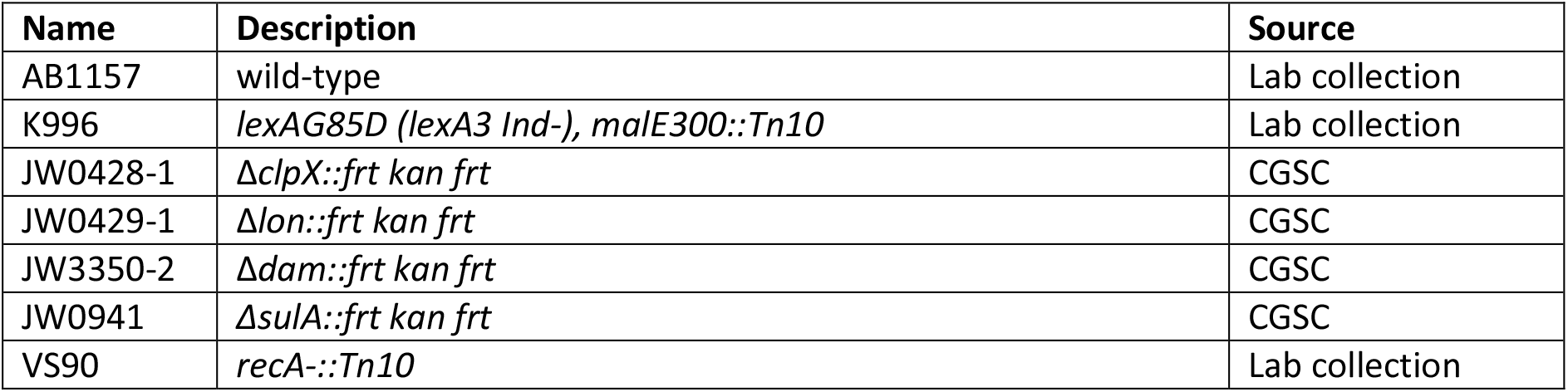
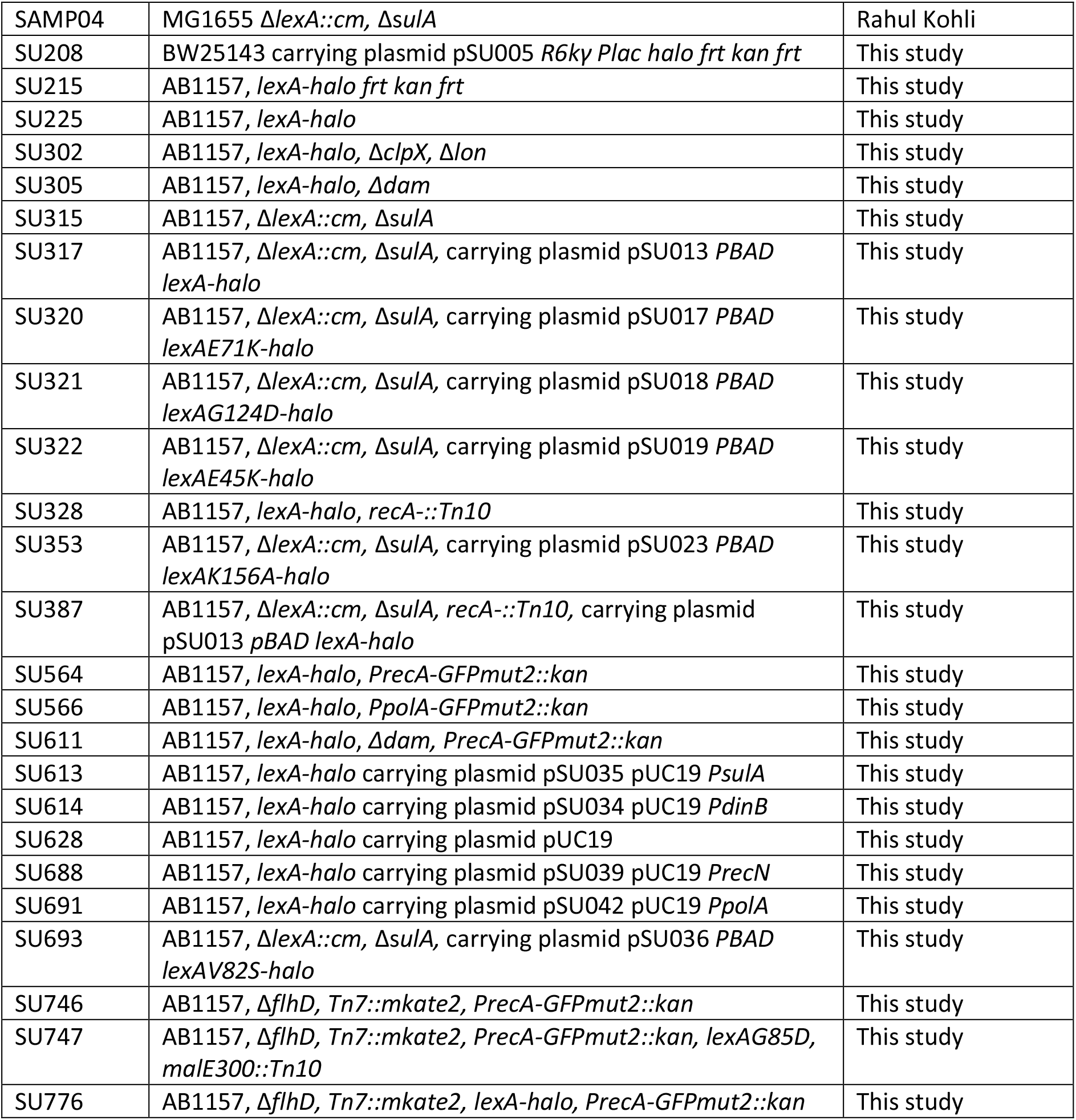

### Cell culture and HaloTag labelling

Strains were streaked from frozen glycerol stocks on to LB agarose with appropriate antibiotic selection. A single colony was used to inoculate LB (+ appropriate antibiotic if strain carried a plasmid) and grown for 6-7 hours. The cultures were then diluted 1:1000 into supplemented M9 minimal medium containing M9 salts (15 g/L KH2PO4, 64 g/L Na2HPO4, 2.5 g/L NaCl, 5.0 g/L NH4Cl), 2 mM MgSO4, 0.1 mM CaCl2, 0.5 μg/ml thiamine, MEM amino acids, 0.1 mg/ml L-proline, 0.2% glucose. Cultures were grown overnight to stationary phase, then diluted 1:100 into supplemented M9 medium and grown to OD600 0.1-0.2 before labelling the HaloTag. For strains expressing LexA-Halo (and LexA mutants) from pBAD plasmid, growth medium containing 0.2% glycerol instead of glucose was used to induce leaky expression. This resulted in similar LexA expression levels as in the strains with chromosomal expression.

We labelled LexA-Halo covalently with TMR dye in live cells according to the protocol described before ^16^. Briefly, cell culture was concentrated 10-fold by centrifugation. 100 μl of cell suspension was incubated with 2.5 μM TMR ligand (Promega) at 25°C for 30 min, followed by 4 rounds of washing, centrifugation, and resuspension in 1 ml of M9 medium to remove free dye from the culture. For dual-colour imaging with PrecA-GFP, we labelled LexA-Halo with JF549 ^40^ instead of TMR, using the same labelling protocol. The enhanced photostability of JF549 compared to TMR increases the number of tracks obtained per cell^16^, which facilitated the single-cell diffusion analysis. After labelling and washing, cells were recovered for 30 min shaking at 37°C to resume growth and allow free dye to diffuse out of cells, which was removed by a further wash with 1 ml of M9 medium. This step was crucial to reduce background fluorescence contamination during single-molecule imaging ^16^. Notably, after the removal of free dye, cell growth and division gradually dilutes the fluorescently-labelled LexA pool as new proteins are synthesised.

For experiments with fixed cells, we labelled LexA-Halo with TMR as above and subsequently resuspended cells in 2.5% paraformaldehyde solution in PBS buffer. Cells were fixed for 30 min at room temperature and washed once in PBS before imaging.

Following labelling, cells were resuspended in approximately 10 μl M9 medium and spotted onto 1% low-fluorescence agarose pads (Bio-Rad) prepared in M9 medium and covered with a no1.5 glass coverslip. Coverslips were treated with air plasma (Plasma Etch) prior to use to remove fluorescent background particles. For UV treatment, cells were placed on the agarose pad and exposed to a UV pulse using a Stratalinker 1800 lamp at the indicated doses before covering with a coverslip. Slides were then incubated at 22°C and multiple fields of view were recorded per slide for up to 45 min for untreated cells, or at the indicated times post UV exposure.

### Single-molecule tracking

Single-molecule imaging was performed using a custom-built total internal reflection fluorescence (TIRF) microscope ^41^ under oblique illumination at 22°C. Initially, cells in a field of view were exposed to 561 nm laser illumination at 0.5 kW cm^-2^ for around 30 seconds such that there was less than one molecule in the fluorescent state per cell on average. We recorded movies of 10-20 thousand frames under continuous 561 nm excitation at 0.5 kW cm^-2^ at a rate of 7.48 ms/frame. Stochastic blinking of TMR or JF549 fluorophores allowed recording hundreds of tracks per cell. A transmitted light image generated by LED illumination was recorded for each field of view for the purpose of cell segmentation.

For dual-colour LexA-Halo and PrecA-GFPmut2 or PpolA-GFPmut2 imaging, we first recorded a 100-ms snapshot with 488 nm illumination at 0.1 kW cm^-2^, followed by a movie of LexA-Halo-JF549 under 561 nm illumination at 0.5 kW cm^-2^ of the same cells.

Custom-written MATLAB software (Mathworks) was used for localisation ^42^, tracking ^43^ and calculation of diffusion coefficients ^44^. Cell outlines were automatically segmented from transmitted light images using a modified version of MicrobeTracker ^45^ combined with SuperSegger ^46^. Apparent diffusion coefficients (D) were calculated from the Mean Squared Displacement (MSD) averaged over 4 steps per molecule: D = MSD/(4·Δt), with Δt = 7.48 ms. Shorter tracks were discarded and longer tracks were truncated to 4 steps. Distributions of diffusion coefficients represent the accumulated tracks of multiple cells in multiple fields of view (as in Fig. 1 & 2) or of single cells (as in Fig. 3).

To quantify the relative abundances of LexA populations (Fig. S2), we fitted a least-squares mixture model to the diffusion coefficient distributions with the sum of the populations constrained to 1. The model curve for the bound population was obtained from measuring LexA-Halo in cells fixed with paraformaldehyde. The model curve for the degraded population was obtained from cells expressing unconjugated HaloTag. The model curve for the free LexA pool was obtained from cells expressing the non-cleavable LexA K156A mutant. Model curves were smoothed using moving mean filters.

For dual-colour measurements, the PrecA-GFPmut2 or PpolA-GFPmut2 intensity per cell was measured from the average pixel intensity within each segmented cell area and the median background intensity outside cells was subtracted.

### In-gel HaloTag fluorescence

Cells expressing HaloTag fusions were grown and labelled with TMR dye as for single-molecule imaging, but washed only once with 1 ml M9 medium. Cells were pelleted after labelling and lysed in 30 μl SDS sample buffer heated at 95°C for 5 min. Halo-tagged species were separated via SDS-PAGE and imaged on a gel scanner with 532 nm laser excitation (Fujifilm Typhoon FLA 7000). Band intensities were quantified using ImageJ and the relative abundance of the degraded LexA population was calculated from the ratio HaloTag/(LexA-Halo + HaloTag).

### Single-cell microfluidics

Mothermachine microfluidic experiments were performed as described^47^ to measure PrecA-GFPmut2 expression dynamics during continuous unperturbed growth in M9 glucose medium at a single-cell level. Cells expressed fluorescent protein mKate2 constitutively and carried an *flhD* gene deletion to remove flagellum motility. Imaging was performed on a Nikon Ti Eclipse inverted fluorescence microscope equipped with perfect focus system, 100x NA1.45 oil immersion objective, sCMOS camera (Hamamatsu Flash 4), motorized stage, and 37°C temperature chamber (Okolabs). Fluorescence images were automatically collected using NIS-Elements software (Nikon) and an LED excitation source (Lumencor SpectraX). Time-lapse movies were recorded at 3-min intervals with 100 ms exposures for GFPmut2 and mKate2 using 50% LED excitation intensities. Movies were analysed using custom Matlab software to segment cells based on cytoplasmic mKate2 fluorescence and to construct single-cell lineages. PrecA-GFPmut2 expression traces represent the average pixel intensity within the area of a cell in each frame after subtracting the median background signal outside cells. Expression pulses were identified by applying a moving mean filter (30 frames window) and using *findpeaks* function in Matlab.

## Acknowledgments

We thank Rahul Kohli (University of Pennsylvania) and David J. Sherratt (University of Oxford) for providing strains, and members of the Uphoff group and David Sherratt’s group for discussions. We thank Aditya Jalin for his contributions during a lab internship. Research in the Uphoff lab is funded by a Wellcome Trust & Royal Society Sir Henry Dale Fellowship (206159/Z/17/Z) and a Wellcome-Beit Prize (206159/Z/17/B). E.C.J. was supported by a Crankstart scholarship. S.U. holds a Hugh Price Fellowship at Jesus College, Oxford. The authors declare no conflict of interest.

