## Supplemental Figures for "Imaging LexA degradation in cells explains regulatory mechanisms and heterogeneity of the SOS response"

### Supplementary Material

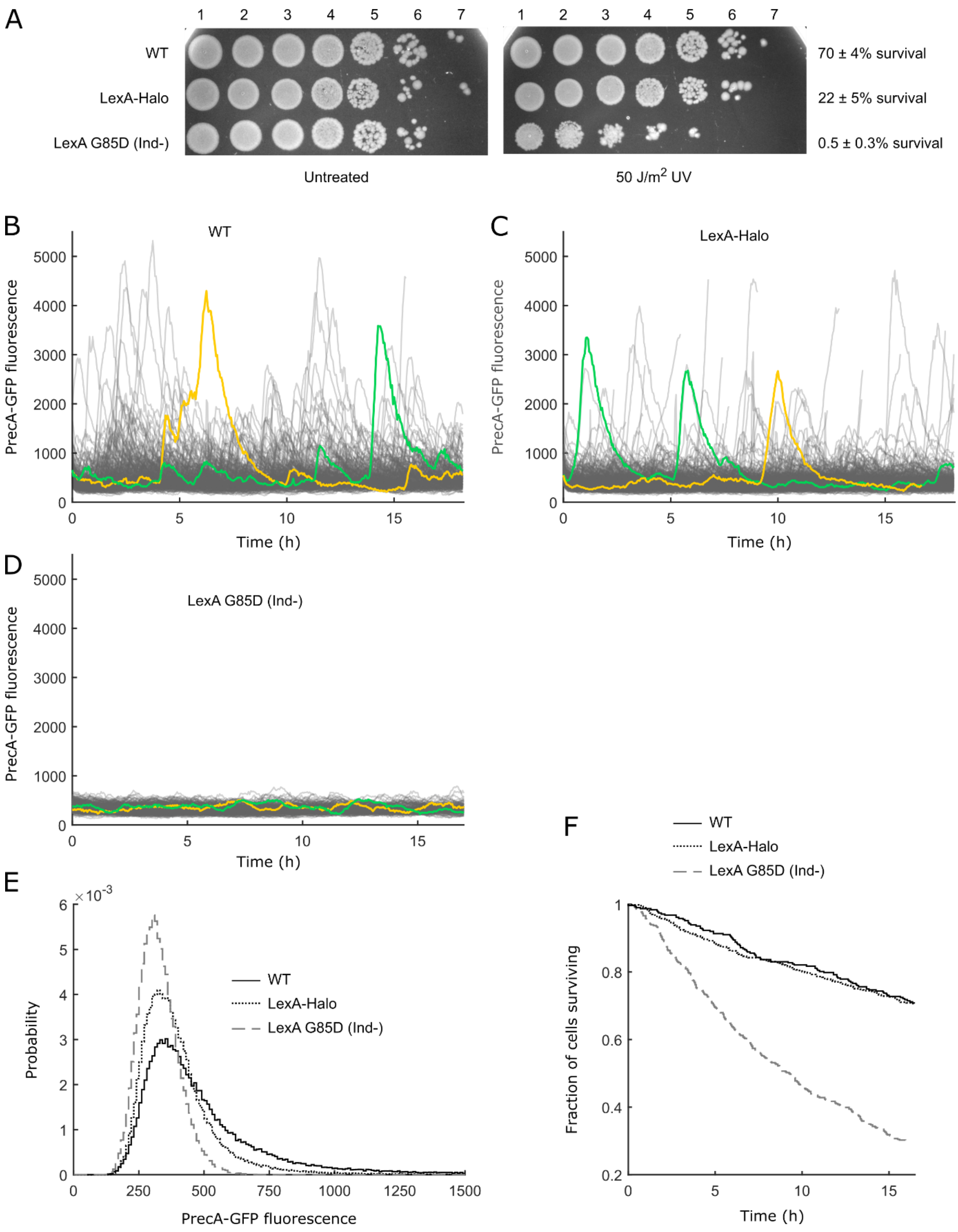


**Fig. S1 Assessing functionality of LexA-Halo fusion. (A)** UV survival assay: 10-fold serial dilutions of over-night LB cultures were spotted on LB agarose plates and exposed to 50 J/m$^2$ UV light before incubating over night. Control plates were incubated without UV exposure. The percentages of surviving cells are the average number of colonies (±SEM) at 10$^6$-fold dilution from 6 independent UV-treated plates divided by the number of colonies on untreated plates. The LexA-Halo fusion strain shows a moderate increase in UV sensitivity compared to the wild-type AB1157 strain, while the non-cleavable LexAG85D mutant is hypersensitive to UV damage. **(B-D)** Single-cell fluorescence dynamics of SOS expression reporter PrecA-GFP during unperturbed growth in mothermachine microfluidic chips for cells expressing wild-type LexA (B, 312 cells), LexA-Halo (C, 501 cells), and non-cleavable LexA G85D (D, 446 cells). Two example cell traces are highlighted per strain. The LexA-Halo fusion strains shows SOS expression pulses similar to the wild-type strain. No pulses are seen in the LexA G85D strain. **(E)** Distributions of PrecA-GFP fluorescence for wild-type LexA, LexA-Halo, and LexA G85D strains show that the LexA-Halo fusion is functional in SOS gene repression. The tail in the distribution for LexA-Halo shows functional SOS induction similar to the wild-type strain. The amplitudes of the SOS expression pulses were slightly reduced compared to the wild-type, matching the reduction in UV tolerance (panel A). **(F)** Distributions of cell survival times during growth in microfluidic chips for wild-type LexA, LexA-Halo, and LexA G85D strains confirms functionality of the LexA-Halo fusion.

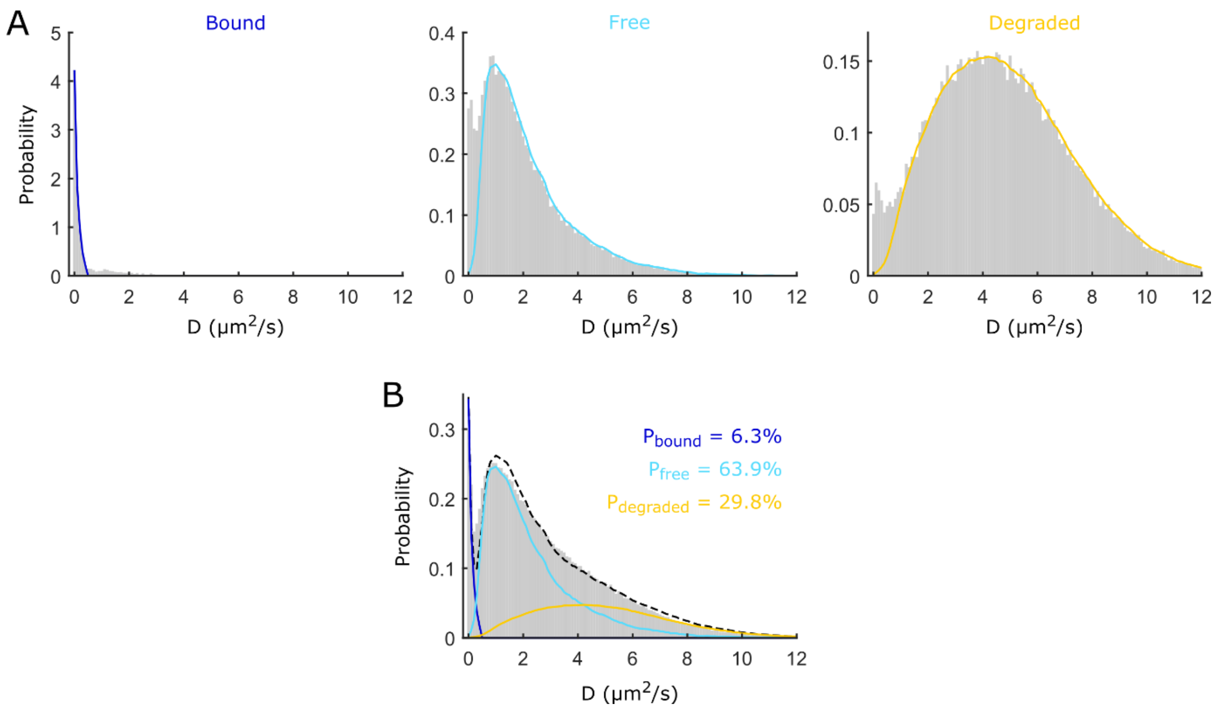


**Fig. S2 Quantifying relative abundances of LexA populations. (A)** Bound, free, and degraded LexA populations were quantified by fitting a mixture model to D distributions using least squares optimisation. The D distribution of LexA-Halo from fixed cells was used as a model for the bound population (dark blue curve). The mobile part of the D distribution for the non-cleavable LexA K156A mutant was used as a model for the free population (light blue curve). The mobile part of the D distribution for the unconjugated HaloTag was used to model the degraded population (yellow curve). The model distributions were smoothed using moving mean filters. **(B)** Fitted mixture model for LexA-Halo in untreated wild-type cells.

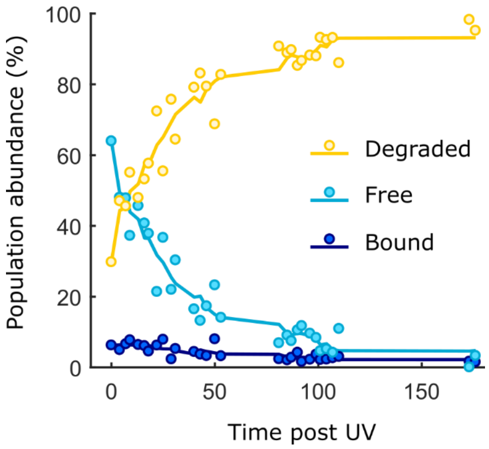


**Fig. S3 Dynamics of LexA populations after UV exposure.** Relative abundances of degraded (yellow), free (light blue), and bound (dark blue) LexA populations after exposing cells to a pulse of 50 J/m$^2$ UV. Bound, free, and degraded LexA populations were quantified by fitting a mixture model to D distributions at different time points post UV using least squares optimisation as shown in Fig. S2. Lines show moving mean curves.

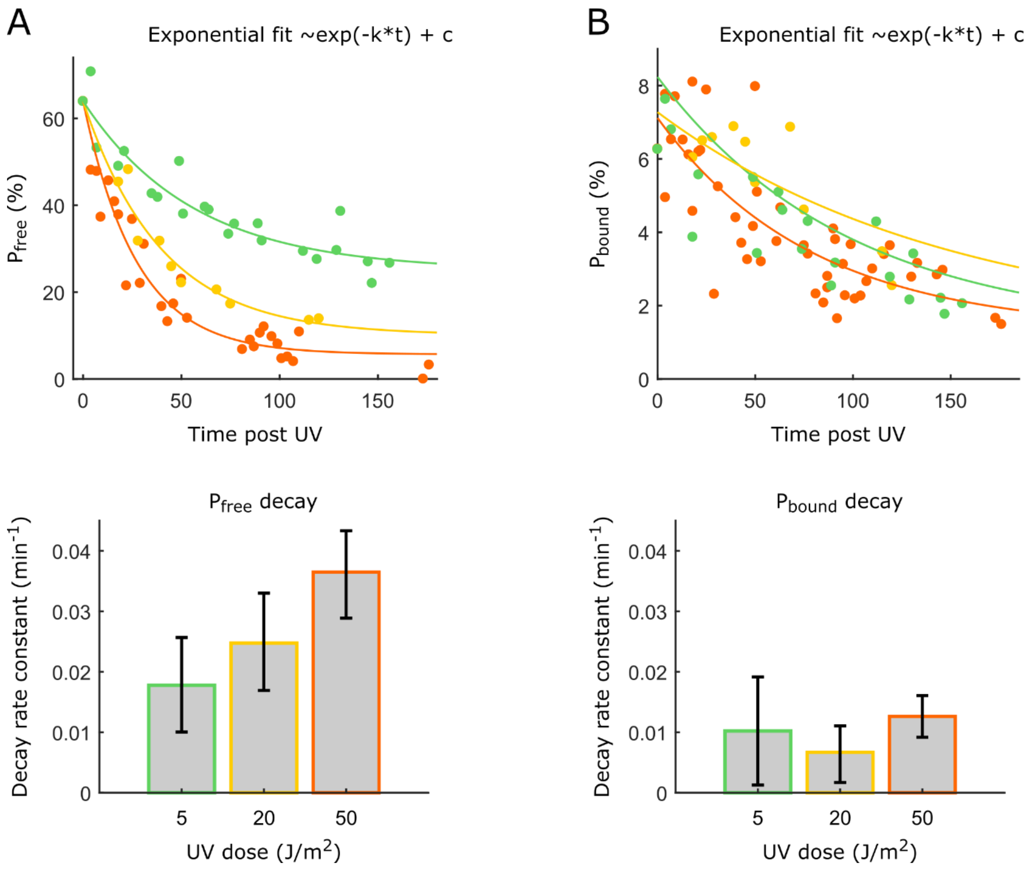


**Fig. S4 Decay of bound and free LexA populations in response to different UV doses.** Decay rate constants were obtained from exponential fits after exposure to 5J/m$^2$ (green), 20J/m$^2$ (yellow), or 50J/m$^2$ (orange). Error bars: 95% confidence intervals. **(A)** The decay rate of the free LexA pool increases with UV dose. **(B)** The decay of the bound LexA population is slower than that of the free LexA population, and the rate does not scale with UV dose.

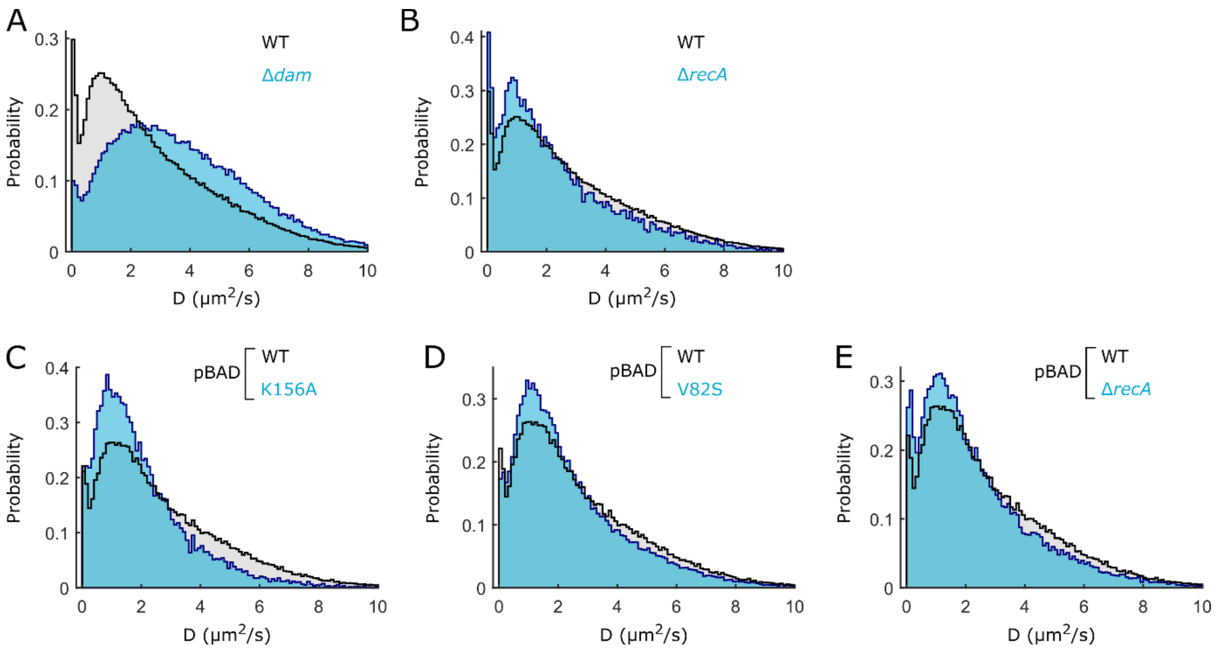


**Fig. S5 D distributions for mutant strains and LexA variants. (A)** LexA-Halo in wild-type (grey, 3146 cells) and Δ*dam* (blue, 837 cells) strain backgrounds. **(B)** LexA-Halo in wild-type (grey, 3146 cells) and *ΔrecA* (blue, 229 cells) strain backgrounds. **(C)** Wild-type LexA-Halo (grey, 838 cells) and non-cleavable LexA K156A mutant (blue, 148 cells) expressed from pBAD plasmid. **(D)** Wild-type LexA-Halo (grey, 838 cells) and auto-cleavage deficient LexA V82S mutant (blue, 582 cells) expressed from pBAD plasmid. **(E)** Wild-type LexA-Halo (grey, 838 cells) expressed from pBAD plasmid in wild-type (grey) and *ΔrecA* (blue, 934 cells) strain backgrounds.

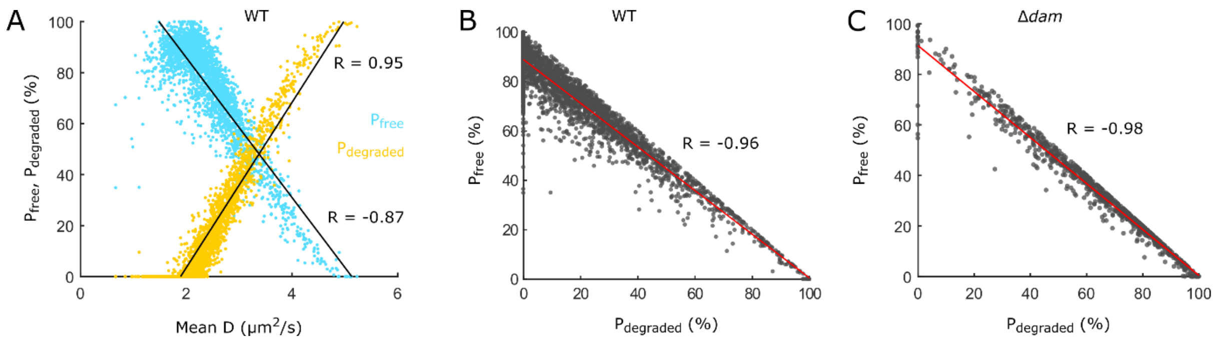


**Fig. S6 Quantification of LexA populations in single cells. (A)** The abundances of $P_{free}$ and $P_{degraded}$ are highly correlated with the average diffusion coefficient of LexA-Halo per cell. Each dot represents a single cell from the wild-type LexA-Halo strain. Black lines show linear fits; R: Pearsson's correlation coefficient. **(B,C)** Plots of $P_{free}$ versus $P_{degraded}$ show that increased level of LexA degradation is correlated with a loss of the pool of free LexA in individual cells from the wild-type and the Δ*dam* strain background. Red lines show linear fits; R: Pearsson's correlation coefficient.

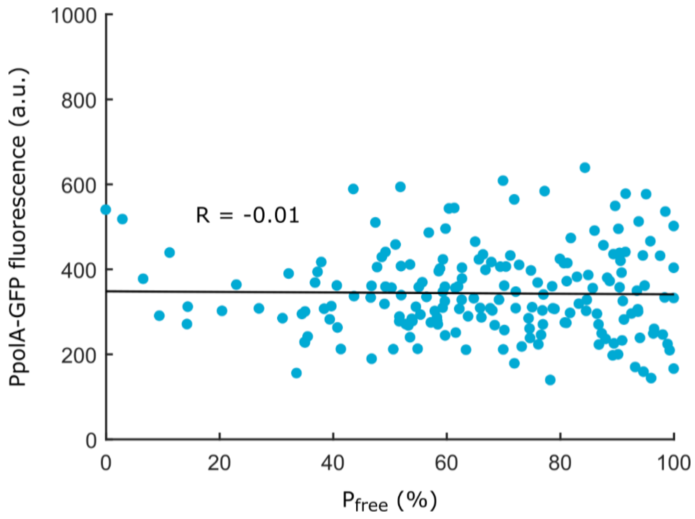


**Fig S7 LexA variability does not correlate with the expression of a gene that is not part of the SOS regulon.** Imaging PpolA-GFP expression and tracking LexA-Halo diffusion in the same cells shows that there is no correlation between gene expression and the free LexA abundance per cell. Each dot represents a single cell. Black line shows linear fit; R: Pearsson's correlation coefficient.
